# STING agonism enhances anti-tumor immune responses and therapeutic efficacy of PARP inhibition in *BRCA*-associated breast cancer

**DOI:** 10.1101/2021.01.26.428337

**Authors:** Constantia Pantelidou, Heta Jadhav, Aditi Kothari, Renyan Liu, Jennifer L. Guerriero, Geoffrey I. Shapiro

## Abstract

Poly (ADP-ribose) polymerase (PARP) inhibitors exert their efficacy by inducing synthetic lethal effects, as well as cGAS/STING-mediated immune responses in BRCA- and other homologous recombination repair-deficient cancer cells. Here we investigated whether the immunologic and therapeutic effects of PARP inhibition in BRCA-deficient breast cancer models could be augmented by synthetic cyclic dinucleotide agonists of STING. Combined PARP inhibition and STING agonism induced a greater degree of STING pathway activation and proinflammatory cytokine production compared to monotherapies in BRCA1-deficient human and mouse triplenegative breast cancer cell lines. In a mouse model of BRCA1-deficient TNBC, the combination also induced an improved immune response compared to either monotherapy alone, evidenced by a greater degree of cytotoxic T cell recruitment and activation, and enhanced dendritic cell activation and antigen presentation. Nanostring mRNA analysis indicated that combinatorial effects were the result of augmented interferon signaling and antigen processing, as well as of heightened leukocyte and dendritic cell functions. Finally, the combination markedly improved anti-tumor efficacy *in vivo* compared to monotherapy treatment, with evidence of complete tumor clearance and prolongation of survival. These results support the development of combined PARP inhibition and STING agonism in *BRCA*-associated breast cancer.

## INTRODUCTION

Inhibitors of poly (ADP-ribose) polymerase (PARP) have improved treatment outcomes of *BRCA*-associated breast cancer and of other homologous recombination (HR) repair-deficient cancers (1). In addition to multiple mechanisms underlying the synthetic lethality of PARP inhibition and HR deficiency (2), PARP inhibitors also induce innate immune responses via cGAS/STING pathway activation, resulting in infiltration of activated cytotoxic T cells, an event critical for maximal efficacy in BRCA-deficient models (3,4). These results have prompted the initiation of clinical trials combining PARP inhibition with immune checkpoint blockade, with the goal of further activating T-cell responses and overcoming PARP inhibitor resistance (5,6). However, preliminary data from these clinical trials have suggested that PD-1/L1 blockade may not enhance efficacy over PARP inhibition alone (6). Recent results have indicated that PARP inhibition may also result in recruitment of immune suppressive macrophages and that PARP inhibitor-mediated T cell activation can be improved with macrophage-targeting strategies (7). Here, we have investigated an alternative approach to improving efficacy in *BRCA*-associated breast cancer by combining PARP inhibition with STING agonism. STING agonists have entered clinical trials and combinations with chemotherapy or immune checkpoint blockade have demonstrated preliminary safety and efficacy (8). In this study, we sought to develop the preclinical rationale for combining STING agonists with PARP inhibition. Our findings demonstrate that the combination of STING agonism and PARP inhibition leads to robust and durable anti-tumor immune responses and complete tumor clearance in *BRCA1*-mutant TNBC models.

## MATERIALS AND METHODS

### Cell Culture

Human MDA-MB-436 cells were purchased from the ATCC and verified with short tandem repeat profiling. Cell line culturing was performed as previously described (3). Cell lines were routinely tested for the presence of mycoplasma using the MycoAlert Mycoplasma Detection Kit (Lonza).

### Compounds

ADU-S100 (MIW815; Chemietek #CT-ADUS100) was reconstituted in USP normal saline (Thermo Fisher Scientific #NC9604723) at 1.25mg/ml. For *in vitro* studies, olaparib (Selleckchem) was reconstituted in DMSO at 100mM. For *in vivo* studies, olaparib (MedChemExpress #HY-10162) was reconstituted in DMSO at 25 mg/ml.

### Immunoblotting

Immunoblotting was performed as previously described (3). The following primary antibodies were used: phospho-TBK1/NAK (Ser172) (D52C2) XP Rabbit mAb [Cell Signaling Technology (CST) #5483S] (for human cells), TBK1 (S172) (Abgent #AP7887a-ev) (for murine cells), TBK1/NAK (D1B4) Rabbit mAb (CST #3504S), phospho-STING (Ser366) (D7C3S) Rabbit mAb (CST #19781S) (for human cells), phospho-STING (Ser365) (D8F4W) Rabbit mAb (CST #72971S) (for murine cells), STING/TMEM173 (Novus Biologicals #NBP224683), Vinculin (CST #4650S).

### Quantitative PCR

RNA was isolated, reverse transcribed and used for quantitative PCR. Primer sequences (5’–3’) for human and murine IFNβ, CCL5 and CXCL10 are previously described (3); murine H2-Aa forward GACCTCCCAGAGACCAGGAT; H2-Aa reverse GGAACACAGTCGCTTGAGGA; murine Clec7a forward CCATAAAAGGCCCAGGGGAT; Clec7a reverse TCGCCAAAATGCTAGGGCA; murine H2-Ab1 forward TGCTACTTCACCAACGGGAC; H2-Ab1 reverse TTTGCTCCAGGCAGACTCAG; murine Ltb forward GATGACAGCAAACCGTCGTG; Ltb reverse CAGCTGTTGAACCCCTGGAT; murine ItgaI forward TGGTCACTGAGCTGTCGTTC; ItgaI reverse CTCAGGATAGGCTGCATGGC.

### *In Vivo* Studies

All animal experiments were conducted in accordance with Institutional Animal Care and Use Committee (approved protocol #17-032). The *K14-Cre;Brca1^f/f^Trp53^f/f^* TNBC mouse model was used as previously described (3). For efficacy studies, treatments were started once tumors reached 150-180 mm^3^ in volume and continued until tumors reached 20 mm in any direction, at which point mice were euthanized. For flow cytometry studies, mice bearing tumors of 150-300 mm^3^ in volume were randomized in treatment groups, so that the average tumor volume in each group was the same. DMSO-reconstituted olaparib was diluted in PBS (Corning) immediately before intraperitoneal injection and administered at 50 mg/kg daily. ADU-S100 was administered intratumorally weekly in a single 40 μl injection of 50 μg, with mice under isoflurane anesthesia. Tumors were measured every 3-4 days using electronic calipers, and tumor volumes were calculated by using the formula (L×W×W)/2.

### Tumor Digestion and Flow Cytometry

At the indicated times, mice were sacrificed, cardiac perfusion was performed and tumors were extracted. A small tumor chunk was snap frozen for RNA analysis and the remainder of the tumor was processed for flow cytometry as previously described (3). The following fluorophore-conjugated primary antibodies were used in flow cytometry studies: Alexa Fluor® 488 antimouse CD45 (BioLegend #103122), Alexa Fluor® 594 anti-mouse CD3 (BioLegend #100240), PE/Cyanine7 anti-mouse CD8a (BioLegend #100721), PE anti-mouse CD4 (BioLegend #100408), Alexa Fluor® 647 anti-human/mouse Granzyme B (BioLegend #515405), Alexa Fluor® 647 mouse IgG1 κ Isotype Ctrl (BioLegend #400135), mouse FoxP3 PerCP/Cy5.5 (BD Biosciences #563902), PerCP/Cy5.5 Rat IgG2a, κ Isotype Ctrl Antibody (BioLegend #400531), Brilliant Violet 605™ anti-T-bet (BioLegend #644817), Brilliant Violet 711™ anti-mouse/human CD11b (BioLegend #101241), Brilliant Violet 650™ anti-mouse CD11c (BD Biosciences #564079), FITC anti-mouse CD40 (BioLegend #124607), Brilliant Violet 421 ™ anti-mouse I-A/I-E (MHCII) (BioLegend #107631). Analysis was performed on FlowJo V10.

### nanoString immune gene expression analysis

RNA isolation from snap-frozen tumor chunks was performed at the Center for Advanced Molecular Diagnostics (CAMD; Brigham and Women’s Hospital) using the Maxwell RSC Tissue kit. RNA was quantified by Nanodrop Spectrophotometer. RNA (100ng) was loaded into the nCounter^®^ PanCancer Immune Profiling Panel, consisting of 770 genes, on the nanoString instrument. Data was analyzed using the Advanced Analysis Module of the nSolver™ analysis software 4.0 (NanoString Technologies) and subjected to Quality control, and Background correction and Normalization against positive controls and housekeeping genes. The geometric mean of eight housekeeping genes was used to calculate normalization factors. Raw counts below the negative controls were discarded from further analysis.

### Statistical Analyses

Statistical analyses were performed using GraphPad Prism v8. For comparison of 2 sets of measurements, unpaired *t*-test was performed. Unpaired *t*-test with Welch correction was used when sample variances were not equal, as defined by the Brown–Forsythe test. For comparison of 3 or more sets of unpaired measurements, one-way ANOVA was performed with Holm-Sidak’s multiple comparisons test on pre-selected relevant pairs. *P* values are indicated on the graphs.

## RESULTS AND DISCUSSION

We previously demonstrated that PARP inhibitors activate the proinflammatory cGAS/STING pathway in BRCA-deficient TNBC cells, eliciting an anti-tumor immune response that is critical for therapeutic efficacy (3). Based on these findings, we hypothesized that pharmacological activation of the cGAS/STING pathway would further augment PARP inhibitor-induced inflammatory signaling and consequent anti-tumor immune responses. We utilized the PARP inhibitor olaparib and the STING agonist ADU-S100, which has been shown to lead to potent anti-tumor immunity in multiple tumor models (9,10) and is currently in clinical trials (8). Immunoblot analysis of phosphorylated STING and its effector TBK1 in the human *BRCA1*-mutant TNBC cell line MDA-MB-436 demonstrated activation of STING-TBK1 signaling in response to olaparib and the higher of the 2 doses of ADU-S100 (**Fig. 1A**, left panel). Combining olaparib with ADU-S100 led to enhanced TBK1 and STING phosphorylation compared to that achieved by the monotherapies, demonstrating increased STING pathway activation. Similarly, in KB1P-G3 cells derived from the *K14-Cre-Brca1^f/f^;Trp53^f/f^* GEMM of TNBC, the Olaparib/ADU-S100 combination led to elevated phospho-TBK1 and phospho-STING levels compared to those observed after single treatments (**Fig. 1A**, right panel).

**Figure 1.**
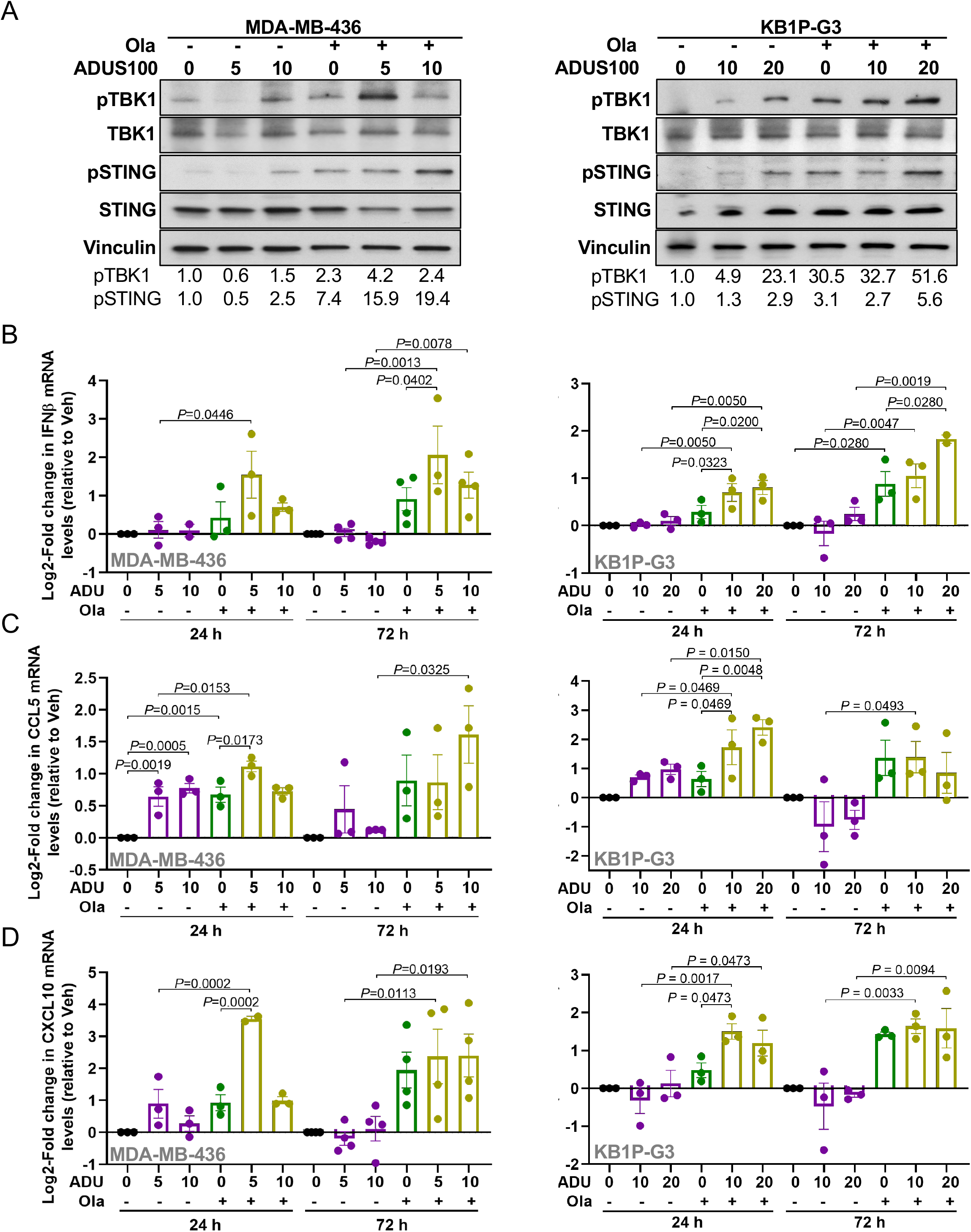
STING agonism enhances STING pathway activation and proinflammatory cytokine production in response to PARP inhibition in TNBC cells. The HR-deficient TNBC cell lines MDA-MB-436 (human) and KB1P-G3 (murine; derived from the K14-*CreBRCA1^f/f^TP53^f/f^* GEMM) were treated with vehicle (-, 0), 1 μM olaparib, the indicated doses of ADU-S100 (in μg/ml) or their combination. **(A)** At 72 hours post-treatment protein was extracted for immunoblot analysis of phospho-TBK1 (Ser172), total TBK1, phospho-STING (Ser366 in human and Ser365 in murine) and total STING expression with vinculin as loading control. Numbers below the blots represent phosphorylated protein levels quantified by densitometric analysis. Immunoblots are representative of 3 independent experiments. **(B-D)** At 24 and 72 hours, post-treatment RNA was extracted and used for qPCR analysis of **(B)** IFNβ, **(C)** CCL5 and **(D)** CXCL10 mRNA expression, plotted as log2-fold change versus vehicle. Error bars represent standard error of the mean (S.E.M.) of 3 independent experiments.

We next examined whether activation of STING signaling resulted in the production of β-interferon (IFNβ) and T cell-attracting chemokines, such as CCL5 and CXCL10. Low-dose ADU-S100 combined with olaparib significantly increased IFNβ mRNA levels in human MDA-MB-436 TNBC cells compared to single treatments, while the higher dose of ADU-S100 did not appear to significantly enhance olaparib-induced IFNβ production (**Fig. 1B**, left panel). In murine cells, the combination of either dose of ADU-S100 with olaparib significantly upregulated IFNβ mRNA levels in comparison to single treatments at 24 hours post-treatment; however, at 72 hours only high-dose ADU-S100 further increased olaparib-induced IFNβ production (**Fig. 1B**, right panel). In MDA-MB-436 cells, the expression of the T-cell attracting chemokines CCL5 and CXCL10 was potently induced by olaparib and ADU-S100 alone but was further increased upon combination of olaparib with low-dose ADU-S100 at 24 hours (**Fig. 1C** and **1D**, left panels). At 72 hours post-treatment however, the increases in CCL5 and CXCL10 production in response to the combination treatment were not statistically significant compared to those achieved by olaparib alone. Similarly, in murine KB1P-G3 cells the combination of ADU-S100 with olaparib resulted in significantly higher CCL5 and CXCL10 mRNA levels compared to those induced by the monotherapies 24 hours after treatment, whereas by 72 hours, olaparib and combination treatments induced comparable CCL5 and CXCL10 production (**Fig. 1C** and **1D** right panels). Of note, at certain doses and time-points, ADU-S100 monotherapy appeared to decrease proinflammatory cytokine production, suggesting the presence of inhibitory feedback loops, as has previously been described after STING pathway activation (3,11). Taken together, our findings indicate that STING agonism enhances PARP inhibitor-induced STING-TBK1 pathway activation and pro-inflammatory cytokine production in human and murine BRCA1-deficient TNBC cells in a dose- and time-dependent manner. Importantly, in human cells, combining lower doses of the STING agonist ADU-S100 with PARP inhibition might lead to more productive inflammatory responses. This observation is consistent with a previous report that higher doses of ADU-S100 can compromise tumor-specific T cell responses and durable antitumor immunity (10).

We next sought to determine whether increased STING-TBK1 pathway activation in response to combined PARP inhibition and STING agonism in cancer cells translated to enhanced antitumor immune responses *in vivo*. To this end, we analyzed *K14-Cre-Brca1^f/f^;Trp53^f/f^* breast tumors implanted in syngeneic mice and treated with olaparib, ADU-S100 or the combination, for the presence and activation of immune cells. Total leukocyte CD45^+^ cell counts were elevated in response to all treatments at 3 (**Fig. 2A**) and 7 days (**Supplementary Fig. S1A, B**). The combination of olaparib and ADU-S100 significantly increased total T-cell counts in comparison to the monotherapies, with both CD8^+^ and CD4^+^ T cell subsets significantly augmented (**Fig. 2A**). Enhanced T-cell recruitment in response to the olaparib/ADU-S100 combination was accompanied by activation of CD8^+^ T cell cytolytic functions, as demonstrated by the significantly increased recruitment of Granzyme-B^+^ CD8^+^ T cells, as well as elevated Granzyme-B total expression (**Fig. 2A**). In the case of CD4^+^ T cells, the combination treatment appeared to increase T-helper 1 and not T-regulatory CD4^+^ T cells, as measured by the expression of their respective markers, Tbet and FoxP3 transcription factors (**Fig. 2A**). Moreover, combining olaparib with ADU-S100 significantly increased the activation and antigen presentation capability of dendritic cells (DCs), as shown by the increases in CD40 and major histocompatibility (MHC) II expression in CD11C^+^CD11B^−^ DCs (**Fig. 2B**). In contrast to the combination therapy that led to a rapid induction of immune responses, olaparib monotherapy-induced T cell recruitment and activation was more evident at 7 days (**Supplementary Fig. S1A**) compared to 3 days post-treatment (**Fig. 2A**). Activated immune cell infiltration following ADU-S100 and olaparib treatment was accompanied by type I IFN production, as evidenced by the significant increase in whole-tumor IFNβ mRNA levels compared to single treatments (**Fig. 2C**). These findings demonstrate that the addition of STING agonism to PARP inhibition elicits a superior immune response compared to either monotherapy alone, characterized by increased cytotoxic T cell recruitment and activation, and enhanced DC activation and antigen presentation.

**Figure 2.**
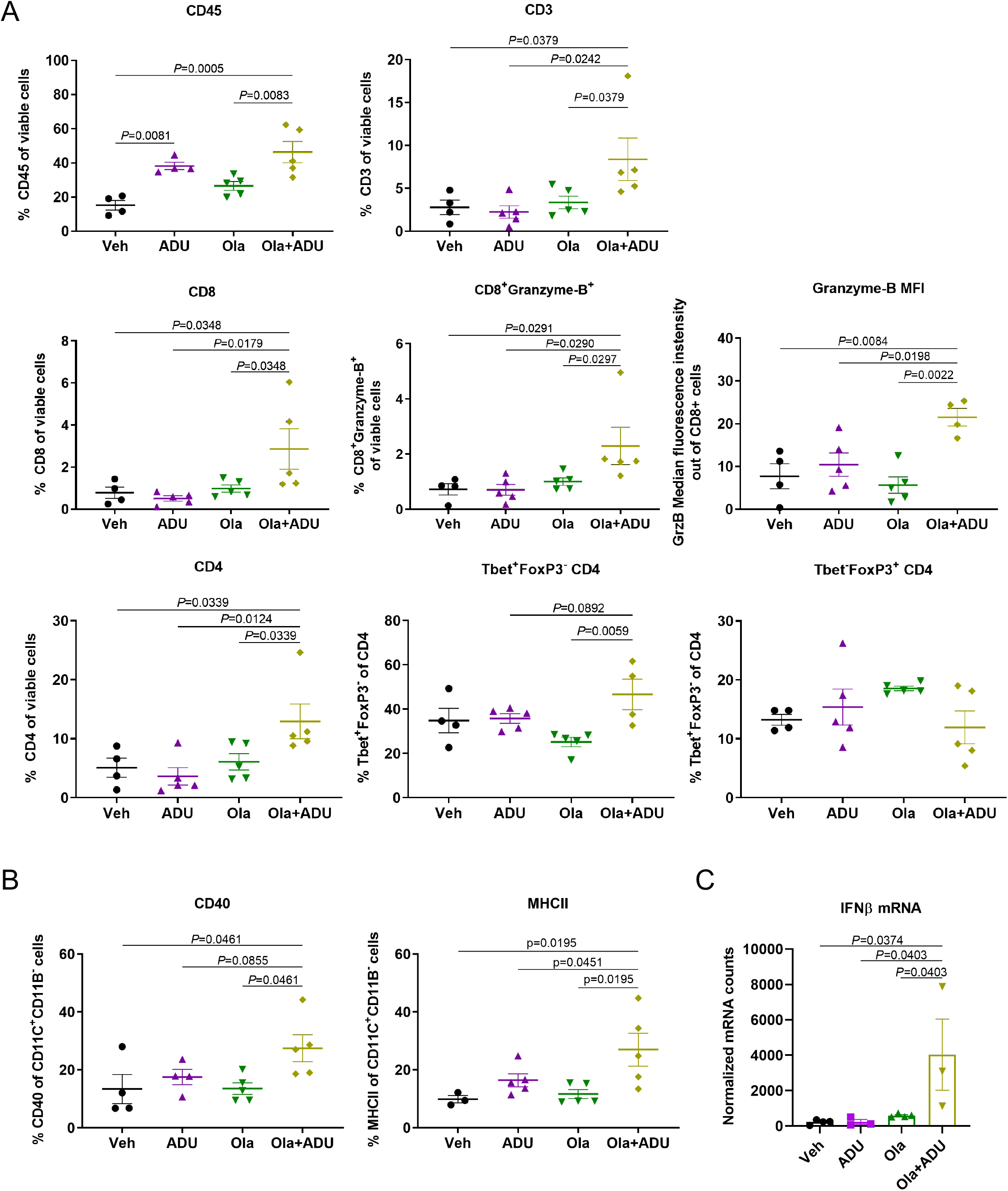
STING agonism and PARP inhibition cooperate to induce anti-tumor immune responses in BRCA-deficient TNBC tumors. Tumor chunks from the *K14-Cre-Brac1^f/f^Tp53^f/f^* GEMM were transplanted in syngeneic FVB/129P mice, which were treated with vehicle, olaparib, ADU-S100 or their combination (4-5 mice/group). **(A, B)** At 3 days, tumors were harvested, and single-cell suspensions were subjected to flow cytometry. Scatter plots show **(A)** CD45^+^ cells, CD3^+^ cells, CD8^+^ and CD4^+^ T-cells, granzyme-B^+^ CD8 T-cells, granzyme-B median fluorescence intensity (MFI) in CD8 cells, Tbet^+^FoxP3^−^ T-helper 1 CD4 cells, Tbet^−^FoxP3^+^ T-regulatory CD4 cells, and **(B)** CD40^+^ and MHCII^+^ CD11C^+^CD11B^−^ dendritic cells. **(C)** At 7 days tumors were harvested, and RNA was isolated and subjected to qPCR analysis of IFNβ mRNA expression. Error bars are S.E.M.

To gain additional mechanistic insights into the anti-tumor immune response induced by the combination of PARP inhibition and STING agonism, we performed nanoString immune gene expression analysis in *K14-Cre-Brca1^f/f^;Trp53^f/f^* tumors treated with olaparib, ADU-S100 or the combination. Gene set analysis revealed that the top-most upregulated genes in response to the olaparib/ADU-S100 combination were involved in antigen processing, MHC, interferon and leukocyte pathways (**Fig. 3A** and **Supplementary Table S1**). The enrichment of these gene sets in response to combination therapy is also illustrated by the nanoString pathway scores (**Supplementary Fig. S2A, B**). In addition to the top-most upregulated gene sets (as compared to vehicle treatment), dendritic cell functions and tumor necrosis factor (TNF) superfamily gene signatures were highly upregulated in comparison to olaparib and ADU-S100 alone (**Fig. 3A** and **Supplementary Table S1**). Differential gene expression is shown by volcano plots (**Fig. 3B** and **Supplementary Fig. S2C)**. Plotting of the normalized mRNA counts confirmed the significant increase in expression of 15 genes in response to the olaparib/ADU-S100 combination as compared to single treatments (**Fig. 3C**). Among the top 5 most significantly induced genes were the H-2 class II histocompatibility antigen, A-B alpha (H2-Aa) and A-K beta (H2-Ab1) chains involved in antigen processing and interferon responses (12); the C-type lectin domain family 7 member A (Clec7a) involved in innate responses, phagocytosis and leucocyte functions (13); the TNF superfamily member Lymphotoxin-beta (Ltb) involved in cytotoxic T-cell effector functions (14); and the integrin alpha L chain (Itgal) involved in leukocyte adhesion and mature T-cell functions (15,16) (**Fig. 3C** and **Supplementary Table S2**). The significant increases in mRNA expression of the top genes were validated by quantitative PCR analysis (**Supplementary Fig. S3A**). In summary, the mRNA expression analysis findings are consistent with the enhanced cytotoxic T cell and DC activation observed by flow cytometry in response to combining STING agonism and PARP inhibition **(Fig. 2** and **Supplementary Fig. S1**). These results demonstrate that the superior anti-tumor immune response observed when STING agonism is combined with PARP inhibition is a result of enhanced interferon signaling, antigen processing, and leukocyte and DC functions.

**Figure 3.**
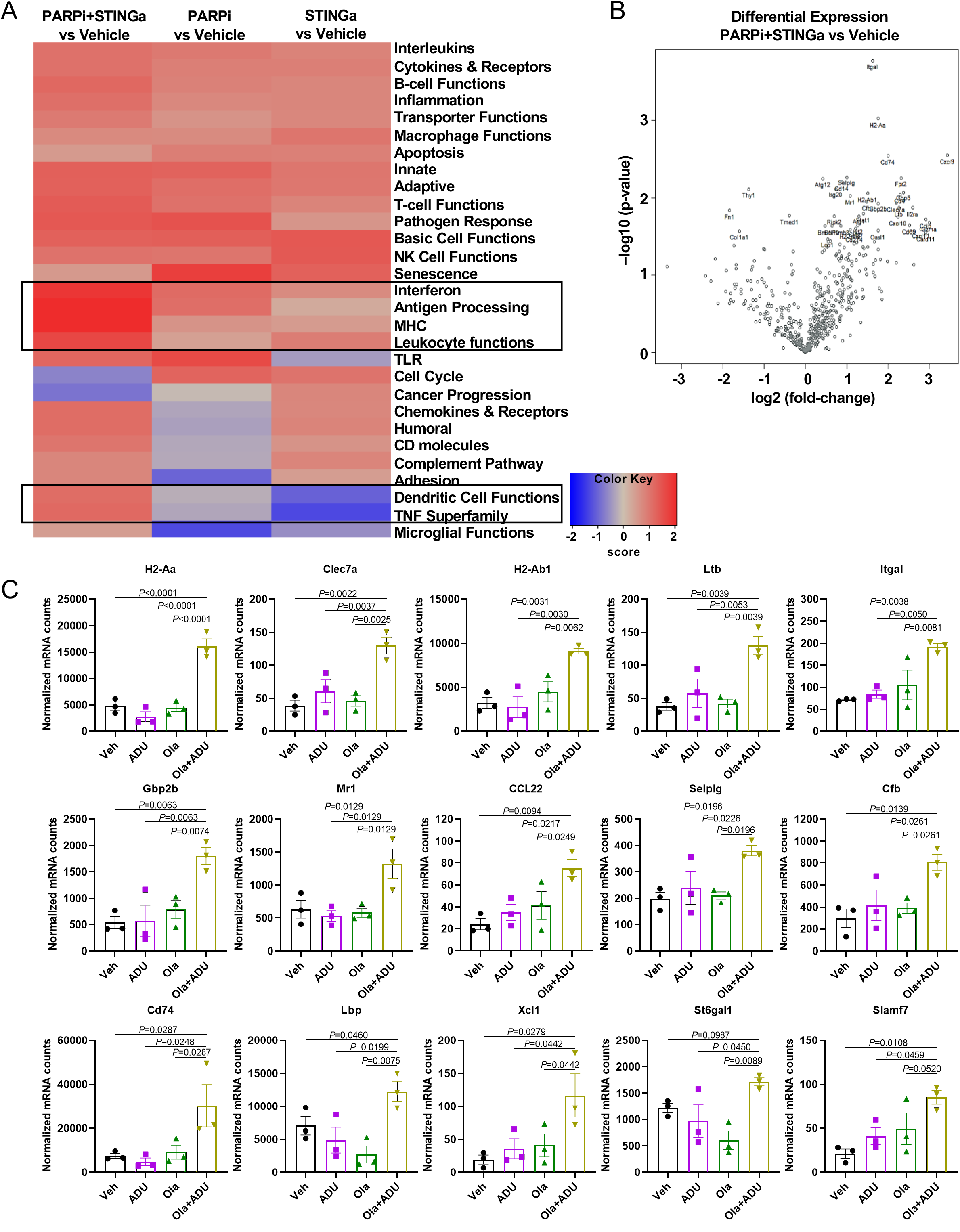
The combination of STING agonism with PARP inhibition upregulates the expression of genes involved in antigen processing, interferon responses and leukocyte functions. *K14-Cre-Brca^f/f^Tp53^f/f^* tumors from mice treated for 3 days with vehicle, olaparib, ADU-S100 or their combination were harvested. RNA was isolated from 3 tumors per group and subjected to nanoString mRNA expression analysis using the nCounter^®^ PanCancer Immune Profiling Panel. **(A)** NanoString Gene Set Analysis (GSA) heatmap of directed global significance scores. The scores measure the extent to which a gene set is up- or down-regulated relative to the covariate and they are calculated as the square root of the mean signed squared t-statistic for the genes in a gene set, with t-statistics coming from the linear regression underlying differential expression analysis. Red denotes gene sets whose genes exhibit extensive over-expression with the covariate, blue denotes gene sets with extensive underexpression. Boxes highlight the most upregulated pathways in response to the combination of PARPi and STINGa. **(B)** Differential expression volcano plot displaying each target’s −log10 (p-value) and log2 fold change in response to the PARPi/STINGa combination treatment versus vehicle treatment. Highly statistically significant targets fall at the top of the plot, and highly differentially expressed genes fall to either side. **(C)** Top 15 most significantly upregulated genes. Scatter plots demonstrate statistically significant increases in normalized RNA counts of indicated genes after treatment with PARPi+STINGa combination in comparison to single treatments. Error bars show S.E.M.

Our observations of augmented immune responses upon combination of a STING agonist with PARP inhibition led us to hypothesize that STING agonism can enhance the therapeutic efficacy of PARP inhibitors in *BRCA*-associated TNBC. We therefore treated mice bearing K14-Cre-*Brca1^f/f^;Tp53^f/f^* tumors with vehicle, olaparib, ADU-S100 or the combination. Treatment with olaparib resulted in tumor shrinkage and a median survival of 129 days (**Fig. 4A-C**). As expected, resistance to olaparib eventually emerged with most tumors relapsing after approximately 100 days of daily treatment (**Fig. 4A**). ADU-S100 alone showed modest therapeutic efficacy and doubled median survival to 27.5 days compared to vehicle control (**Fig. 4A, C**). The combination of ADU-S100 with olaparib resulted in significantly greater reduction in tumor volume than olaparib as early as week 1 (day 7), with most notable differences observed after week 8 (day 56) (**Fig. 4A**, **B**). Remarkably, the combination treatment led to complete tumor clearance in all enrolled mice (**Fig. 4A)** and 100% tumor-free survival (**Fig. 4C**). All treatments were well tolerated, and no animal weight loss was observed after long-term exposure (**Fig. 4D**). These findings demonstrate that STING agonism maximizes the antitumor efficacy of PARP inhibition and overcomes PARP inhibitor resistance in *BRCA*-associated TNBC models.

**Figure 4.**
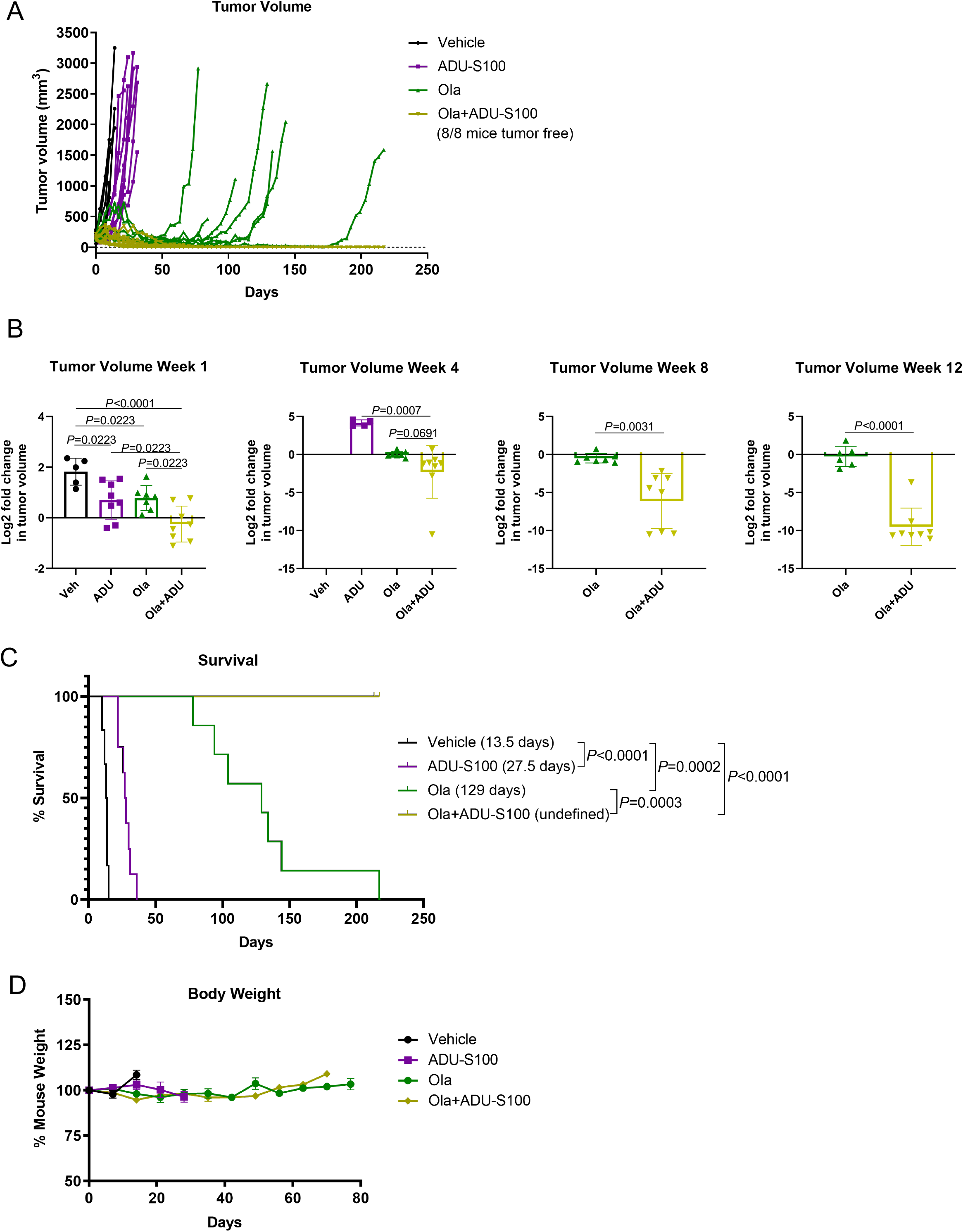
STING agonism potentiates the therapeutic efficacy of PARP inhibitors in BRCA-deficient models of TNBC and overcomes resistance. Tumor chunks from the K14-*Cre-Brca1^f/f^Tp53^f/f^* GEMM were transplanted in syngeneic FVB/129P mice, which were treated with vehicle, olaparib, ADU-S100 or their combination (6-8 mice/group). Tumor volume was measured twice a week and survival was recorded. **(A)** Tumor volumes in individual mice over time. **(B)** Log2 fold change in tumor volumes at week 1, 4, 8, 12. **(C)** Percent survival. Median survival shown in brackets. Statistical analysis was performed using the Log-rank (Mantel-Cox) test (*P* values indicated). Note: Intratumoral ADU-S100 injections stopped when the tumor was cleared. **(D)** Weight of mice treated with vehicle, olaparib, ADU-S100 or their combination long term. Plot demonstrates that the combination treatment is well tolerated.

PARP inhibitor-induced STING pathway activation and anti-tumor immune responses have been demonstrated in multiple tumor models, providing rationale for exploiting combinations of PARP inhibitors with immunotherapies for improved therapeutic efficacy (17,18). This work has demonstrated that intratumoral activation of the proinflammatory STING pathway by PARP inhibitors is amplified by synthetic cyclic dinucleotide agonists of STING, leading to robust and durable anti-tumor immune responses in *BRCA1*-mutant TNBC models. It will be important to extend these findings to other tumor models and to determine whether similar results can be achieved with systemic agonists of STING, currently under development (19,20). In conclusion, the potent preclinical therapeutic efficacy of combined PARP inhibition and STING agonism warrants further development of this regimen as a treatment for *BRCA*-associated TNBC.

## Notes

**Financial Support**: This work was supported by the Dana-Farber/Harvard Cancer Center Specialized Program of Research Excellence (SPORE) in Breast Cancer, NIH grant P50 CA 168504 (G.I.S.), a SPORE Career Enhancement Award (J.L.G.), the Ludwig Center for Cancer Research at Harvard (J.L.G., G.I.S.) and the Emerson Foundation (C.P., G.I.S.).

**Conflict of Interest Statement:** G.I.S. has received research funding from Eli Lilly, Merck KGaA/EMD-Serono, Merck, and Sierra Oncology. He has served on advisory boards for Pfizer, Eli Lilly, G1 Therapeutics, Roche, Merck KGaA/EMD-Serono, Sierra Oncology, Bicycle Therapeutics, Fusion Pharmaceuticals, Cybrexa Therapeutics, Astex, Almac, Ipsen, Bayer, Angiex, Daiichi Sankyo, Seattle Genetics, Boehringer Ingelheim, ImmunoMet, Asana, Artios, Atrin, Concarlo Holdings, Syros, Zentalis and CytomX Therapeutics. In addition, he holds a patent entitled, “Dosage regimen for sapacitabine and seliciclib,” also issued to Cyclacel Pharmaceuticals, and a pending patent, entitled, “Compositions and Methods for Predicting Response and Resistance to CDK4/6 Inhibition,” together with Liam Cornell. J.L.G. is a consultant for GlaxoSmithKline (GSK), Codagenix, Verseau, Kymera and Array BioPharma and receives sponsored research support from GSK, Array BioPharma and Eli Lilly.

### Competing Interest Statement

G.I.S. has received research funding from Eli Lilly, Merck KGaA/EMD-Serono, Merck, and Sierra Oncology. He has served on advisory boards for Pfizer, Eli Lilly, G1 Therapeutics, Roche, Merck KGaA/EMD-Serono, Sierra Oncology, Bicycle Therapeutics, Fusion Pharmaceuticals, Cybrexa Therapeutics, Astex, Almac, Ipsen, Bayer, Angiex, Daiichi Sankyo, Seattle Genetics, Boehringer Ingelheim, ImmunoMet, Asana, Artios, Atrin, Concarlo Holdings, Syros, Zentalis and CytomX Therapeutics. In addition, he holds a patent entitled, Dosage regimen for sapacitabine and seliciclib, also issued to Cyclacel Pharmaceuticals, and a pending patent, entitled, Compositions and Methods for Predicting Response and Resistance to CDK4/6 Inhibition, together with Liam Cornell. J.L.G. is a consultant for GlaxoSmithKline (GSK), Codagenix, Verseau, Kymera and Array BioPharma and receives sponsored research support from GSK, Array BioPharma and Eli Lilly.

