## Supplementary Figures and Tables for "STING agonism enhances anti-tumor immune responses and therapeutic efficacy of PARP inhibition in *BRCA*-associated breast cancer"

### SUPPLEMENTARY FIGURE LEGENDS

**Supplementary Figure S1. Immunologic effects of combined PARP inhibition and STING agonism in a BRCA-deficient GEMM at 7 days.** (A) Tumor chunks from the K14-*Cre-Brca1*$^{f/f}$*Tp53*$^{f/f}$ GEMM were transplanted in syngeneic FVB/129P mice, which were treated with vehicle, olaparib, ADU-S100 or their combination (4-5 mice/group). At 7 days tumors were harvested, and single-cell suspensions were subjected to flow cytometry. Scatter plots show CD45$^{+}$ cells, CD3$^{+}$ cells, CD8$^{+}$ and CD4$^{+}$ T-cells, granzyme-B$^{+}$ CD8 T-cells, granzyme-B median fluorescence intensity (MFI) in CD8 cells, and CD40$^{+}$ CD11C$^{+}$CD11B$^{-}$ dendritic cells. Error bars are S.E.M. (B) Gating strategy used in the flow cytometric analysis of immune cell subsets. Debris was excluded on SSC vs FSC plot and zombie aqua-negative, i.e., viable cells were gated. Live cells were analyzed for expression of CD45 (hematopoietic cells), CD3 (total T cells), CD8 T-cells, CD4 T-cells and dual expression of CD11B/CD11C. CD8$^{+}$ T-cells were further analyzed for expression of Granzyme-B and median fluorescence intensity (MFI) was derived. CD4 T-cells were analyzed for expression of Tbet and FoxP3. CD11C$^{+}$CD11B$^{-}$ dendritic cells were analyzed for expression of CD40 and MHCII.

**Supplementary Figure S2. Additional nanostring analyses of tumors from mice treated with PARP inhibition and STING agonism.** K14-*CreBRCA1*$^{f/f}$*TP53*$^{f/f}$ tumors from mice treated for 3 days with vehicle, olaparib, ADU-S100 or their combination were harvested. RNA was isolated from 3 tumors per group and subjected to nanoString mRNA expression analysis using the nCounter® PanCancer Immune Profiling Panel. (A) Quality control heatmap of the normalized data, scaled to give all genes equal variance, generated via unsupervised clustering. Orange indicates high expression; blue indicates low expression. Each treatment group has 3 samples. (B) Violin plot of the nanoString pathway scores in the top 3 upregulated pathways in response to STINGa+PARPi. The Pathway Score summarizes the data from a pathway's genes with a single score. (C) Volcano plot of the differentially expressed genes in response to the PARPi/STINGa combination treatment versus STINGa (left panel) or PARPi (right panel) single treatment.

**Supplementary Figure S3. Validation of the top upregulated genes from nanoString analysis by qPCR.** Scatter plots show significant increases in the RNA levels of H2-Aa, Clec7a, H2-Ab1, Itgal and Ltb genes (top 5 in the nanoString mRNA analysis) in response to the combination treatment as measured by qPCR.

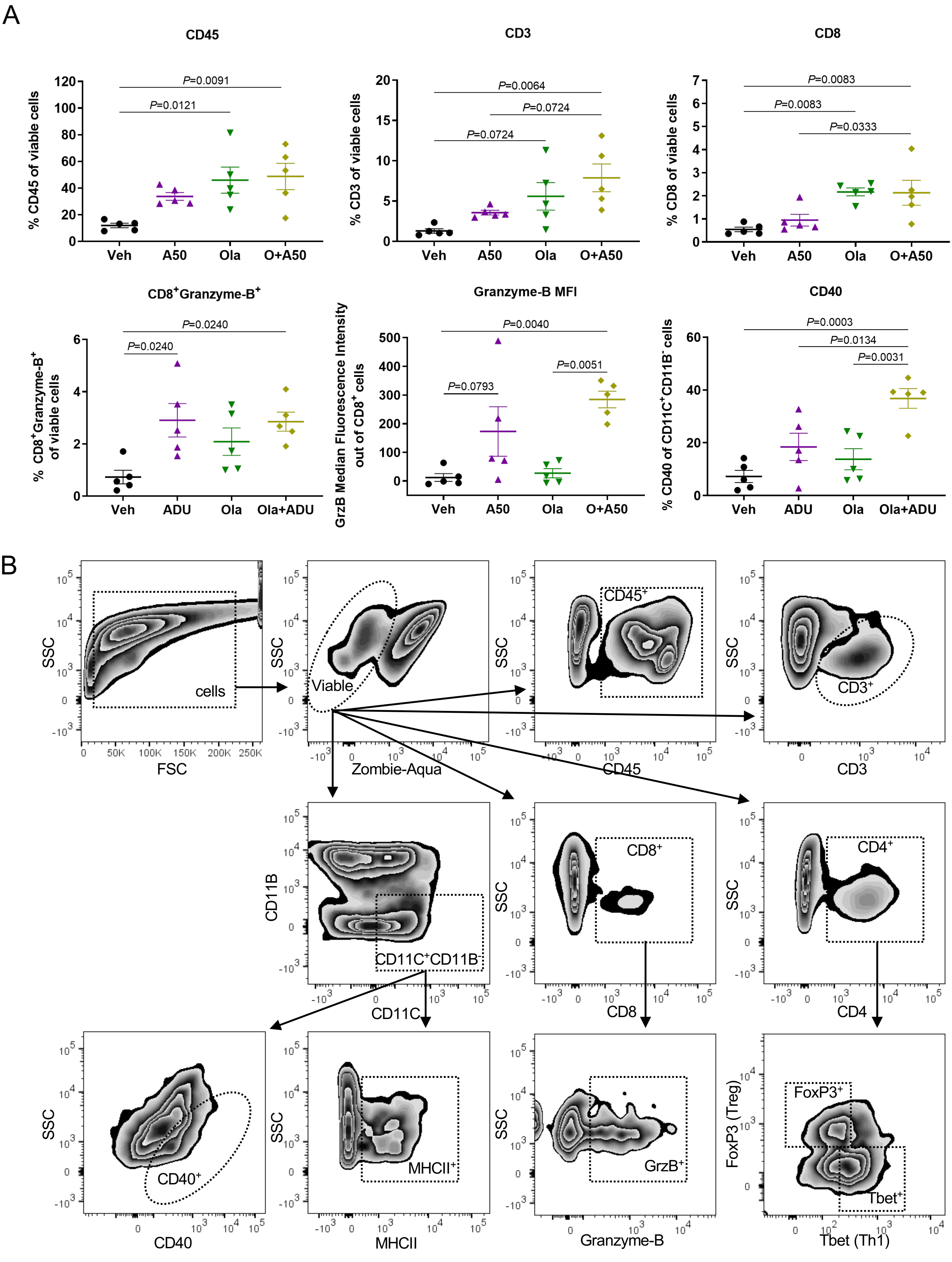


Supplementary Fig. S1

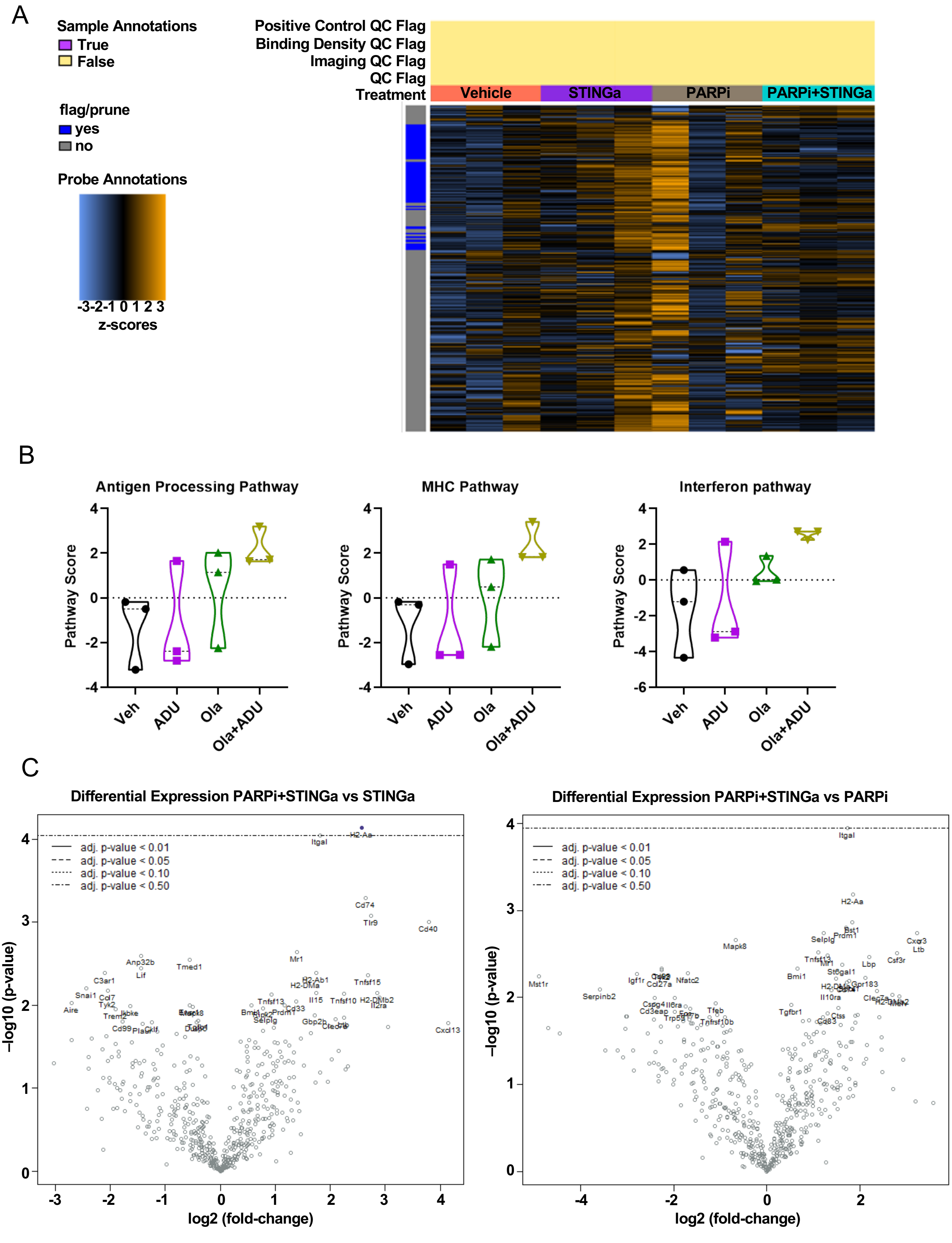


Supplementary Fig. S2

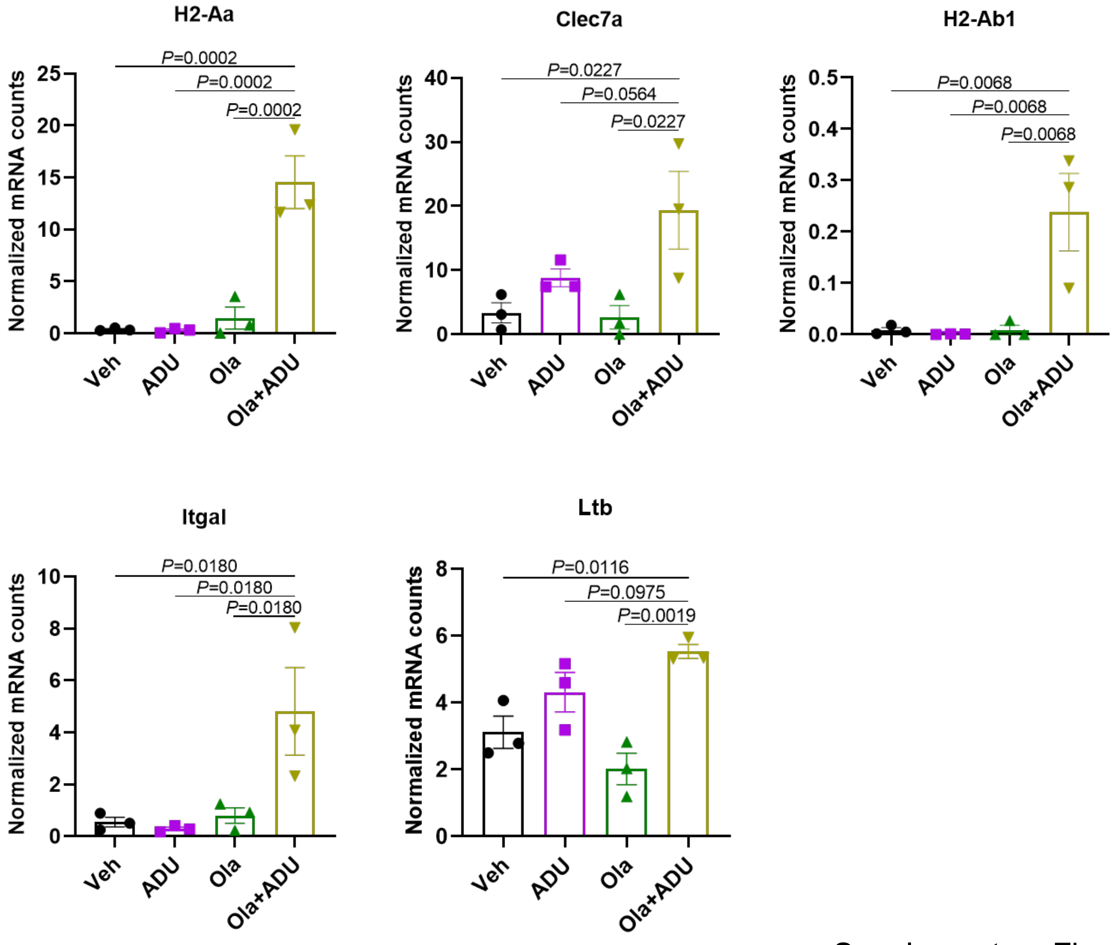


Supplementary Fig. S3

**<u>Supplementary Table S1</u>: Directed Global Significance Scores - differential expression in each treatment group vs baseline of vehicle**

| Gene Set | Olaparib+ADU-S100 | Olaparib | ADU-S100 |
|---|---|---|---|
| Antigen Processing | 2.084 | 1.14 | 0.295 |
| MHC | 2.075 | 0.625 | 0.512 |
| Interferon | 1.922 | 1.18 | 0.748 |
| Leukocyte Functions | 1.69 | 0.442 | 0.939 |
| Pathogen Response | 1.457 | 1.54 | 0.602 |
| T-Cell Functions | 1.414 | 1.095 | 0.834 |
| Innate | 1.337 | 1.296 | 1.034 |
| Basic Cell Functions | 1.325 | 1.355 | 1.457 |
| Adaptive | 1.32 | 1.145 | 1.02 |
| TLR | 1.269 | 1.656 | -0.393 |
| B-Cell Functions | 1.231 | 0.874 | 0.807 |
| TNF Superfamily | 1.212 | -0.288 | -1.291 |
| Humoral | 1.169 | -0.352 | 0.842 |
| Chemokines & Receptors | 1.166 | -0.295 | 0.796 |
| Dendritic Cell Functions | 1.156 | -0.202 | -1.015 |
| Inflammation | 1.137 | 0.768 | 0.805 |
| Cytokines & Receptors | 1.106 | 0.903 | 0.904 |
| Interleukins | 1.106 | 0.981 | 0.854 |
| NK Cell Functions | 1.081 | 1.044 | 1.445 |
| CD molecules | 1.044 | -0.277 | 0.631 |
| Transporter Functions | 0.97 | 0.626 | 0.786 |
| Complement Pathway | 0.808 | -0.256 | 0.822 |
| Adhesion | 0.801 | -0.984 | 0.478 |
| Macrophage Functions | 0.752 | 0.734 | 1.042 |
| Senescence | 0.552 | 1.8 | 1.324 |
| Apoptosis | 0.535 | 0.916 | 0.883 |
| Microglial Functions | 0.485 | -1.278 | -0.546 |
| Cell Cycle | -0.659 | 1.275 | 1.073 |
| Cancer Progression | -0.815 | -0.072 | 0.804 |

**Supplementary Table S2: Annotation of top upregulated genes identified by nanoString analysis in response to STINGa+PARPi treatment**

| nCounter® Mouse PanCancer Immune Profiling Panel - Annotations | | | |
|---|---|---|---|
| # | Gene Name | Protein | Annotation |
| 1 | *H2-Aa* | H-2 class II histocompatibility antigen, A-B alpha chain | Antigen Processing, Interferon Response, Mature T-Cell Functions, MHC class I & II |
| 2 | *Clec7a* | C-type lectin domain family 7 member A | Inflammatory Response, Innate Response, Leukocyte Functions, Phagocytosis |
| 3 | *H2-Ab1* | H-2 class II histocompatibility antigen, A-K beta chain | Antigen Processing, Interferon Response, Mature T-Cell Functions, MHC class I & II |
| 4 | *Ltb* | Lymphotoxin-beta | Cytokines, Interleukins, TNF Superfamily Members |
| 5 | *Itgal* | Integrin alpha L | Adhesion, CD molecules, Leukocyte Functions, Mature T-Cell Functions |
| 6 | *Gbp2b* | Interferon-induced guanylate-binding protein 1 | Basic Cell Functions |
| 7 | *Mr1* | Major histocompatibility complex class I-related gene protein | Antigen Processing, MHC class I & II |
| 8 | *Ccl22* | C-C motif chemokine 22 | Chemokines, Cytokines, Humoral Response, Inflammatory Response, Regulation of Inflammatory Response |
| 9 | *Selplg* | P-selectin glycoprotein ligand 1 | Adhesion, CD molecules, Leukocyte Functions |
| 10 | *Cfb* | Complement factor B | Complement Pathway, Innate Response |
| 11 | *Cd74* | Cluster of Differentiation 74 (HLA class II histocompatibility antigen gamma chain) | Antigen Processing, CD molecules, Innate Response, Mature B-Cell Functions, MHC class I & II, T-Cell Differentiation |
| 12 | *Lbp* | Lipopolysaccharide-binding protein | Chemokines, Inflammatory Response, Innate Response, Leukocyte Functions, Macrophage Functions, Transmembrane Transportation |
| 13 | *Xcl1* | Lymphotactin | Chemokines, Cytokines, Cytotoxicity, Inflammatory Response, Innate Response, Mature T-Cell Functions |
| 14 | *St6gal1* | Beta-galactoside alpha-2,6-sialyltransferase 1 | Basic Cell Functions, Humoral Response |
| 15 | *Slamf7* | Signaling lymphocytic activation molecule (SLAM) family member 7 | CD molecules, Innate Response, Natural Killer Cell Functions |
